# SNAP23 is required for constitutive and regulated exocytosis in mouse oocytes

**DOI:** 10.1101/562843

**Authors:** Lisa M. Mehlmann, Tracy F. Uliasz, Katie M. Lowther

## Abstract

Mammalian oocytes are stored in the ovary for prolonged periods, arrested in meiotic prophase. During this period, their plasma membranes are constantly being recycled by endocytosis and exocytosis. However, the function of this membrane turnover is unknown. Here, we investigated the requirement for exocytosis in the maintenance of meiotic arrest. Using Trim-away, a newly developed method for rapidly and specifically depleting proteins in oocytes, we have identified the SNARE protein, SNAP23, to be required for meiotic arrest. Degradation of SNAP23 causes premature meiotic resumption in follicle-enclosed oocytes. The reduction in SNAP23 is associated with loss of gap junction communication between the oocyte and surrounding follicle cells. Reduction of SNAP23 protein also inhibits regulated exocytosis in response to a Ca^2+^ stimulus (cortical granule exocytosis), as measured by lectin staining and cleavage of ZP2. Our results show an essential role for SNAP23 in two key processes that occur in mouse oocytes and eggs.

**Summary:** The SNARE protein, SNAP23, is required to maintain gap junction communication between the oocyte and follicle cells that is needed to maintain oocyte meiotic arrest, as well as for cortical granule exocytosis at fertilization.

## INTRODUCTION

Mammalian oocytes arrest at prophase of the first meiosis for an extended period of time, during which they accumulate RNAs and proteins that enable them to sustain the development of an early embryo. During this period of meiotic arrest, they constitutively secrete factors such as GDF9, BMP15, and FGF8, which are critical for proper follicle development [1, 2]. In return, the follicle cells provide the oocyte with substances needed for protein synthesis and metabolism, cholesterol, and importantly, the cGMP that is necessary for the maintenance of meiotic arrest prior to the LH surge [2–5]. Prior to the LH surge, oocytes undergo robust endocytosis that probably plays roles in the uptake of extracellular materials. Endocytosis also seems to be involved in cell signaling during meiotic arrest, as blocking receptor-mediated endocytosis in mouse oocytes prevents oocyte maturation by elevating cAMP levels [6]. In addition, constitutive exocytosis in the oocyte may be required to replenish lipids and membrane proteins that are taken up during constitutive endocytosis.

During meiotic maturation, the oocyte undergoes numerous cytoplasmic changes that are required for successful embryonic development after fertilization [7]. One change that occurs is the development of the ability to undergo Ca^2+^-regulated exocytosis in the form of cortical granule exocytosis. Cortical granules are vesicles that reside in the cortical region of the mature egg opposite the meiotic spindle [8, 9], and their exocytosis occurs in response to a large release of intracellular Ca^2+^ at fertilization [10]. Upon exocytosis, the enzyme ovastacin, contained in the granules, cleaves the zona pellucida protein ZP2 and renders the zona pellucida impermeable to additional sperm [11]. Thus, cortical granule exocytosis is an important polyspermy preventing mechanism. Interestingly, immature mouse oocytes are unable to undergo cortical granule exocytosis even when stimulated by Ca^2+^ [10, 12], so the ability to undergo exocytosis develops during maturation.

The proteins required for exocytosis have not been well studied in mammalian oocytes or eggs. (For simplicity, we will refer to immature, prophase-arrested oocytes herein as “oocytes,” and mature, metaphase II-stage oocytes as “eggs”). It is likely that the process is similar to exocytic events that occur in other types of secretory cells. For example, intracellular Ca^2+^ triggers exocytosis of neurotransmitters in neurons [13–15], and insulin receptor activation causes exocytosis of the GLUT4 transporter to the plasma membrane in adipocytes [16, 17]. These exocytic events are mediated by SNARE (soluble *N*-ethylmaleimide-sensitive factor attachment protein receptor) proteins in which two t-SNARES (associated with the target membrane) form a complex with one v-SNARE (associated with the vesicle destined for fusion; Fig. 1). The t-SNAREs are a syntaxin and either SNAP25 or SNAP23 and the v-SNARE is a vesicle-associated membrane protein (VAMP, also called synaptobrevin). SNARE proteins contain coiled coil domains that form a very stable, parallel 4 helix bundle. One helix is contributed by syntaxin, one by VAMP, and two by SNAP25/SNAP23. Formation of the SNARE complex allows the vesicle and target membranes to come into close contact and fuse, releasing the contents of the vesicles to the extracellular space.

**Figure 1.**
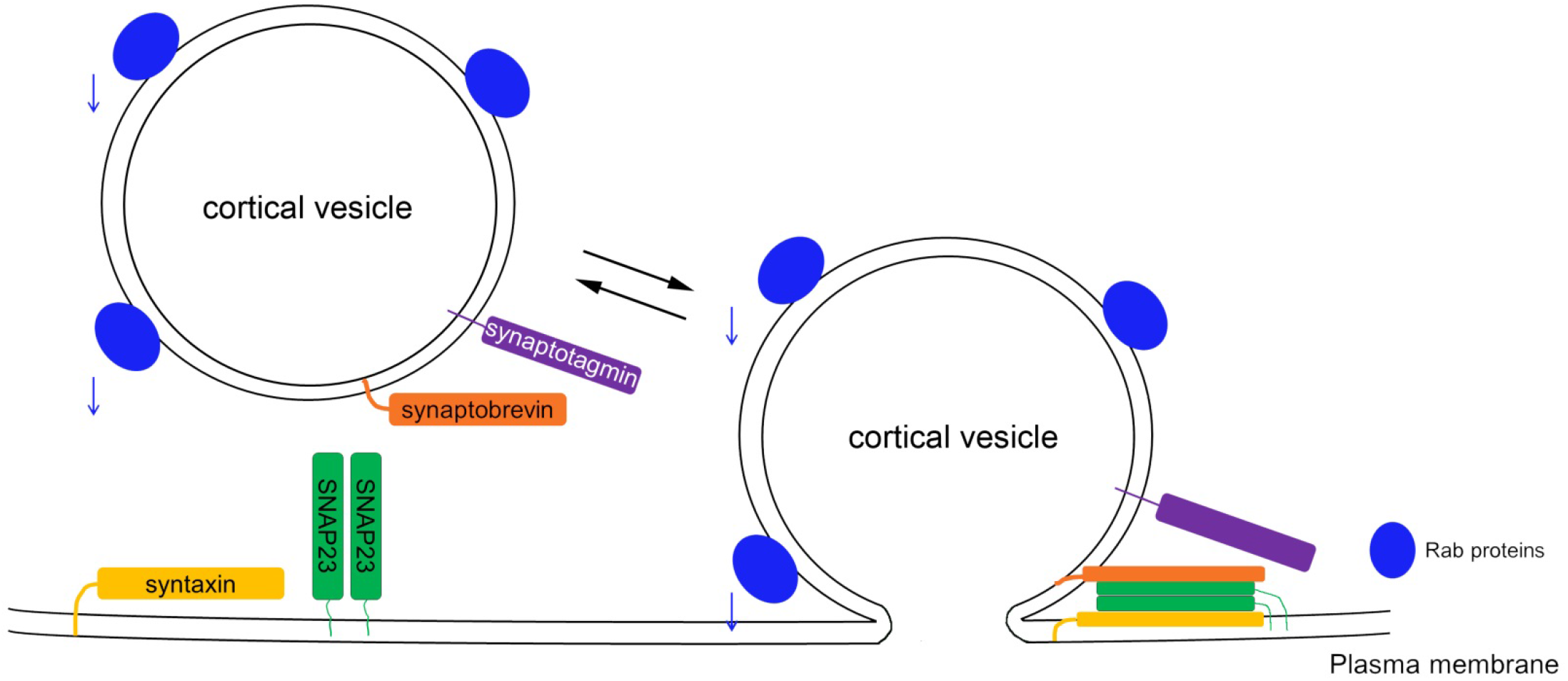
SNARE proteins that may be involved in exocytosis in mouse oocytes and eggs. Coiled coil domains from t-SNAREs (syntaxin and SNAP23/SNAP25) zip together with a v-SNARE (synaptobrevin) to pull the vesicle membrane close to the plasma membrane and facilitate membrane fusion.

Regulated exocytosis has been studied to some extent in eggs of a variety of species. Sea urchin eggs contain homologs of mammalian secretory proteins [18], and inhibition of the SNARE synaptobrevin (VAMP) inhibits cortical granule exocytosis [19]. In porcine eggs, cortical granules associate with SNAP23, syntaxin 2, and VAMP1 [20], but the function of these SNARE proteins has not been examined in detail. It was reported previously that SNAP25 is required for regulated exocytosis in mouse eggs [21], based on the finding that botulinum toxin, which cleaves SNAP25 [22], inhibits cortical granule exocytosis. Syntaxin 4 was identified in mouse eggs, but no function was investigated [23]. Other SNARE proteins have not been identified, but proteins in the Rab family have been implicated in cortical granule exocytosis. In particular, Rabs 3A and 27A are of interest based on their co-localization with cortical granules [24], and on the findings that microinjection of function-blocking antibodies [24] or protein depletion using RNAi [25] inhibits cortical granule exocytosis [24, 25]. In addition to Rab proteins, other key players in the SNARE pathway, α-SNAP and NSF (N-ethylmaleimide sensitive factor) [26] appear to affect cortical granule exocytosis. In this study, we investigated the role of the SNARE protein, SNAP23, in constitutive and regulated exocytosis in mouse oocytes and eggs, respectively. We used a newly developed method, “Trim-away,” to identify SNAP23 as essential for both of these processes.

## MATERIALS AND METHODS

### Media and Reagents

Except where noted, all chemicals were obtained from Millipore Sigma (St. Louis, MO). The medium used to collect oocytes was MEMα (Gibco 12000022, Thermo Fisher, Waltham, MA), supplemented with 20 mM HEPES, 75 μg/ml penicillin G, 50 μg/ml streptomycin, 0.1% polyvinyl alcohol (PVA), and 10 μM milrinone to inhibit spontaneous meiotic resumption. For in vitro maturation, oocytes were transferred to bicarbonate-buffered MEMα, in which the HEPES was replaced with 25 mM sodium bicarbonate and the PVA was replaced with 5% fetal bovine serum (Invitrogen). Media were equilibrated with 5% CO_2_ and 95% air. The medium used for follicle dissection and microinjection was bicarbonate-buffered MEMα supplemented with 3 mg/ml bovine serum albumin, 5 μg/ml insulin, 5 μg/ml transferrin, 5 ng/ml selenium, and 10 ng/ml ovine follicle stimulating hormone (FSH; provided by A.F. Parlow, National Hormone and Peptide Program, Torrance, CA).

### DNA constructs and antibodies

Fluorescently tagged TRIM21 constructs (TRIM21-GFP and TRIM21-RFP) were obtained from Melina Schuh (Max Planck Institute for Biophysical Chemistry; [27]). To make mRNA, the constructs were linearized with XbaI and in vitro transcribed and polyadenylated using the mMessage mMachine T7 Ultra kit (Thermo Fisher). Wild type (SNAP25WT) and dominant negative (SNAP25Δ20) SNAP25 constructs were obtained from Khaled Machaca (Weill Cornell Medicine Qatar) and mRNA was synthesized after linearizing the plasmids with NheI. Approximately 10 pg of mRNA was injected into oocytes. Human SNAP23-GFP, cloned into the Sgfl-Mlu1 site of the pCMV6-AC-GFP vector, was purchased from OriGene (catalog #RG201596) and was subcloned into the SmaI site of the pIVT vector (provided by Carmen Williams, NIEHS)[28] after digesting the plasmid with Pme1 and Kpn1, followed by T4 DNA polymerase treatment to blunt the Kpn1 end. mRNA was synthesized with the mMessage mMachine T7 kit after linearizing the plasmid with NdeI. Approximately 3 pg of mRNA was injected into oocytes for SNAP23-GFP expression.

The following antibodies were used: ZP2 was obtained from Jurrien Dean (NIDDK) [29]; SNAP25 was from BD Biosciences (San Jose, CA) #610366; SNAP23 used for the blot in Fig. 2 was from Synaptic Systems (Göttingen, Germany) #110 203, and for all other experiments was from Fisher (#PA1-738); and affinity purified antibody against the IP_3_ receptor was provided by James Watras (UConn Health). Rabbit IgG was from Sigma (#I5006). Horseradish peroxidase-conjugated secondary antibodies were from Santa Cruz (Santa Cruz, CA) or Advansta Inc. (Menlo Park, CA). Two other SNAP25 antibodies were used to test for SNAP25 expression in oocytes and eggs, which yielded identical results as the antibody from BD Biosciences. These were Santa Cruz sc-58289 and Cell Signaling Technology #5308.

**Figure 2.**
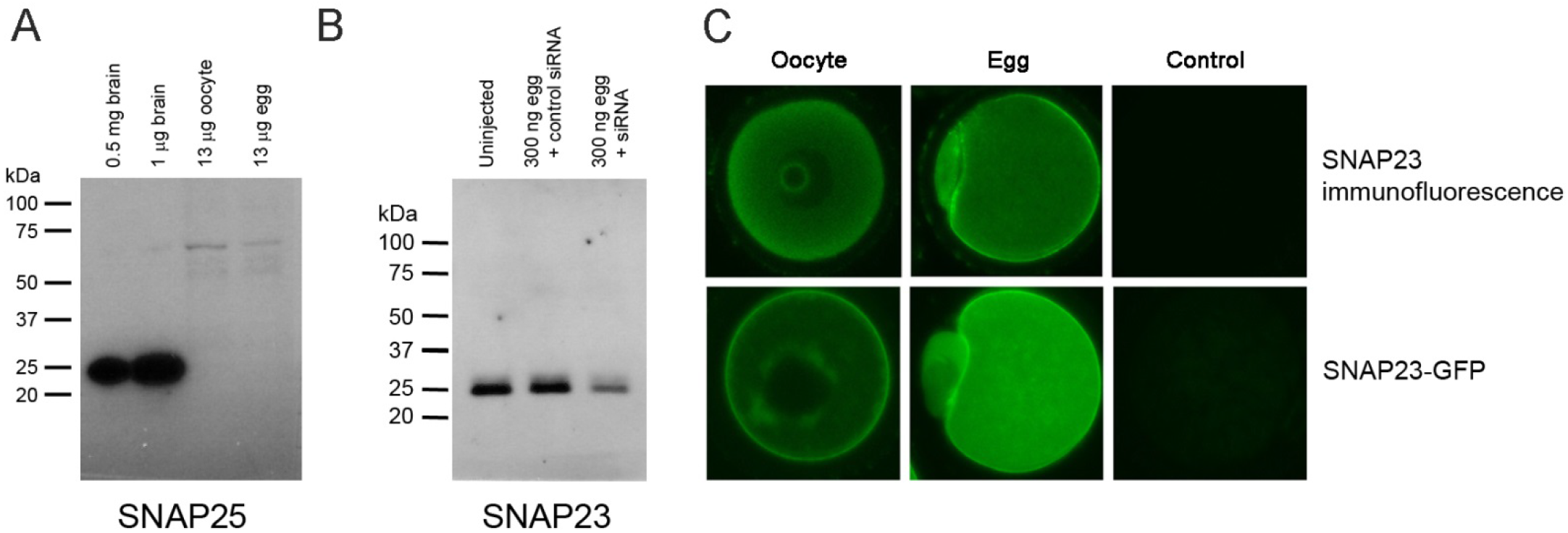
SNAP23, but not SNAP25, is expressed in mouse oocytes and eggs. A) Western blot showing SNAP25 expression in brain, but not in mouse oocytes or eggs; lysates from 525 oocytes or eggs (13 μg total protein) were used per lane. B) SNAP23 is expressed in mouse eggs. Blot shows lysates from 12 isolated oocytes that were not injected; injected with scrambled siRNA; or injected with siRNA targeting SNAP23 following a ~29 hr incubation and overnight maturation to produce mature eggs. C) Localization of SNAP23 in oocytes and eggs. Top panels are immunofluorescence and bottom panels are SNAP23-GFP expression. Controls = secondary antibody only (top) or uninjected egg (bottom).

### Mouse follicle and oocyte isolation, culture, and microinjection

All experiments were done with prior approval of the Animal Care and Use Committee at UConn Health. Preantral or antral stage follicles that were ~140-180 μm or ≥290 μm in diameter, respectively, were manually dissected from ovaries of 23-26-day-old B6SJLF1 mice (Jackson Laboratories, Bar Harbor, ME) as described previously [30]. Following isolation, follicles were plated on Millicell culture plate inserts (PICMORG50, Millipore, Billerica, MA) and were cultured in a humidified atmosphere at 37° with 5% CO_2_, 95% air. After a 3 hr culture period, follicle-enclosed oocytes were examined under an upright microscope for the presence of a germinal vesicle (GV). Only follicles containing oocytes with readily visible GVs were selected for use in these experiments. Fully grown, GV-stage oocytes were obtained from the ovaries of 6-12-week-old NSA (CF1) mice (Envigo, South Easton, MA). Cumulus cells were removed using a small-bore pipet.

Quantitative microinjection of follicle-enclosed oocytes was performed as described previously [30]. Four-5 follicles were loaded into a microinjection chamber between two coverslips and follicles were removed and re-plated onto Millicells following each set of injections. In some cases, follicles were removed from Millicells, reloaded into microinjection chambers, and the oocytes injected with 10 pl of 1 mM Alexa488 (Thermo Fisher), diluted in PBS (50 μM final concentration in the oocyte). Dye transfer from the oocyte to the surrounding follicle cells was visualized using a 20X, 0.5 NA lens on a Zeiss Pascal confocal microscope. Isolated oocytes were injected as described previously [6, 31]. Final injected concentrations were calculated based on an oocyte volume of 200 pl.

### Trim-away

Oocytes were injected with a mixture of mRNA encoding fluorescently tagged TRIM21 (10 pl of 0.3 μg/μl) and SNAP23 antibody or purified IgG (50 μg/ml final injected amount). Prior to injection, antibody/IgG was spin-dialyzed into PBS and concentrated using either Amicon Ultra - 0.5 ml, 10 kDa centrifugal filters (Millipore) or Vivaspin 500 50 kDa cutoff filters (GE Healthcare). Concentrated antibody/IgG solutions were diluted into PBS containing 0.05% NP-40 to reduce stickiness [27, 32].

### Western blotting

Oocytes and eggs were counted and washed into HEPES-buffered MEMα to remove serum, then were transferred to 0.5 ml microcentrifuge tubes in a small volume of medium (1-2 μl). 15 μl of 1X Laemmli sample buffer was added to each tube, and samples were stored at −20° until use. Total protein was estimated using a value of 25 ng protein per oocyte [33]. Western blotting was performed as previously described [34]. Proteins were separated in 4-20% polyacrylamide gels (Bio-Rad, Hercules, CA). Electrophoresed proteins were transferred to nitrocellulose or PVDF membranes and were blocked either in TBST containing 5% nonfat dry milk (NFDM) or 5% BSA, or in Advansta block (Advansta). SNAP25 antibody was diluted in 5% NFDM-containing TBST to a concentration of 0.25 μg/ml and SNAP23 antibody was diluted in Advansta block to a concentration of 2 μg/ml. Both of these antibodies were incubated on blots overnight at 4°C. ZP2 blots were blocked overnight in 5% BSA-containing TBST, and the antibody, diluted 1:1000 in blocking buffer, was incubated on the blots for 2 hrs at RT. Blots were then washed and incubated with horseradish peroxidase-conjugated secondary antibodies and were developed using either ECL Prime (GE Healthcare, Piscataway, NJ) or WesternBright Sirius horseradish peroxidase substrate (Advansta), using a charge-coupled device camera (G:box Chemi XT4; Syngene (Frederick, MD).

### Immunofluorescence and lectin staining

Oocytes and eggs were fixed for 30 min in PBS containing 0.1% PVA (PBS-PVA) and 2% formaldehyde, permeabilized for 15 min in PBS-PVA and 0.1% Triton X-100, and blocked for at least 15 min in PBS containing 3% BSA and 0.01% Triton X-100. Oocytes and eggs were incubated in primary antibody, diluted to 10 μg/ml in blocking buffer, overnight at 4°C. After washing in PBS-PVA, eggs were incubated in Alexa488-conjugated secondary antibody for 1-2 hrs at room temperature. Oocytes and eggs were washed in PBS-PVA, then were imaged using a Zeiss Pascal confocal microscope using a 40X 1.2 NA water immersion objective (C-Apochromat; Carl Zeiss MicroImaging, Inc., Thornwood, NY).

To label cortical granules, proteins were depleted using Trim-away as outlined above. After RNA/antibody injections, zonae pellucidae were removed with 5 mg/ml pronase in HEPES-buffered MEMα. Oocytes were matured overnight in bicarbonate buffered MEMα with 5% FBS. After maturation, eggs were imaged to confirm that they were fluorescent, demonstrating that the TRIM21 protein was expressed. Eggs were then treated with 100 μM thimerosal (ICN Biochemicals, Cleveland, OH) for 45 min in HEPES-buffered MEMα to stimulate intracellular Ca^2+^ oscillations [35], fixed for 30 min with 2% formaldehyde in PBS-PVA, blocked 30 min in PBS containing 3 mg/ml BSA and 0.1 M glycine, permeabilized 20 min in PBS-PVA containing 0.1% Triton X-100, washed 3 times in PBS-PVA, incubated 30 min in 50 μg/ml TRITC-Lens culinaris agglutinin (LCA) in PBS-PVA, washed 3 times in PBS-PVA, then imaged with the 40X 1.2 NA lens on the confocal microscope.

### Ca^2+^ imaging

SNAP23 was depleted in oocytes using Trim-away (using RFP-tagged TRIM21) and injected oocytes were matured overnight. Uninjected eggs were used as a control. Mature eggs were loaded with Fluo3-AM (Molecular Probes) for 30 min per the manufacturer’s instructions, using PowerLoad to aid in the uptake of the dye and probenecid to inhibit intracellular compartmentalization. After dye loading, eggs were adhered to glass coverslips in 90 μl drops of HEPES buffered medium containing no PVA. Five-6 eggs were imaged simultaneously using a 40X, 1.2 NA lens on the confocal microscope. After obtaining a basal Ca^2+^ level, Ca^2+^ was recorded at one scan per second during addition of 100 μM thimerosal (10 μl of 1 mM stock to ensure rapid mixing). Following recordings, a region of interest was measured across a stacked image in ImageJ, values were entered into Excel, and results are presented as the amplitude of each measurement (*F*) subtracted by the baseline (*F*_0_) and divided by the baseline signal [(*F*-*F*_0_)/*F*_0_].

### Statistical Analysis

GraphPad Prism was used for statistical analyses. Fisher’s exact test was used to determine statistical significance of categorical data and Student’s t-test was used to determine statistical significance of quantitative data. P<0.05 was considered to be signicant.

## RESULTS AND DISCUSSION

### SNAP23, but not SNAP25, protein is detectable in mouse oocytes and eggs

A previous study reported that botulinum neurotoxin A (BoNT/A) significantly inhibits cortical granule exocytosis when injected into mouse eggs [21]. Based on this observation, and on western blots showing expression of SNAP25 in eggs, the authors concluded that SNAP25, which is mainly expressed throughout the nervous system [36, 37], is essential for cortical granule exocytosis. However, BoNT/A can also cleave the closely related SNARE, SNAP23, in vitro [38] and based on the high degree of similarity between SNAP25 and SNAP23, it is possible that antibodies might crossreact. To investigate which of these proteins are expressed in mouse oocytes and eggs, we first tried to confirm expression of SNAP25 using western blot. We were unable to detect SNAP25 protein in oocytes or eggs, using 3 different antibodies against SNAP25 and relatively high amounts of oocyte and egg lysates (Fig. 2A and Fig. S1). These antibodies were different than the one used previously [21], and they all showed identical results. In contrast, the closely related and ubiquitously expressed SNAP23 was abundantly expressed (Fig. 2B and Fig. S1). We confirmed the identity of SNAP23 by targeting the mRNA encoding it by microinjection of siRNA. The SNAP23 siRNA reduced SNAP23 protein to 44% of the control after a 29 hr incubation following microinjection (Fig. 2B).

We localized SNAP23 in oocytes and eggs using immunofluorescence and a SNAP23-GFP fusion protein. In oocytes, both methods showed the protein mainly concentrated at the egg cortex, with some interior staining and staining around the nucleolus or the GV (Fig. 2C). In eggs, both immunofluorescence and SNAP23-GFP showed a thin band of SNAP23 around the entire egg cortex, with some interior staining as well. The SNAP23-GFP eggs had higher background fluorescence than the eggs stained with immunofluorescence, which could be due to overexpression of the protein. Generally there was staining above the meiotic spindle in eggs, but in some cases there was a region near the spindle that had less staining. This area was much smaller than the cortical granule-free area that occurs in the vicinity of the meiotic spindle (see Fig. 5). These results demonstrate that SNAP23, rather than SNAP25, is expressed in oocytes and eggs and show that it is located at an appropriate region of the oocyte/egg to be involved in exocytosis.

### Depleting SNAP23 stimulates meiotic resumption in follicle-enclosed oocytes

To test a role for SNAP23 in constitutive exocytosis, we depleted it in follicle-enclosed oocytes. Depletion of SNAP23 in oocytes using siRNA was incomplete, as levels were only reduced by ~56% after a two day culture period (Fig. 2B). Therefore, we depleted SNAP23 using a newly developed method called “Trim-away” [27]. This method utilizes TRIM21, an intracellular antibody receptor and E3 ubiquitin ligase. When TRIM21 is expressed in oocytes or eggs followed by injection of a specific antibody against the protein of interest, TRIM21 binds to the antibody and recruits the protein/antibody complex to the proteasome, causing rapid degradation [39]. Using this method, we found that SNAP23 protein was reduced to a nearly undetectable level within ~3 hrs after injecting SNAP23 antibody into TRIM21-expressing oocytes (Fig. 3A). The TRIM21 protein was required for SNAP23 protein depletion, as microinjection of SNAP23 antibody alone did not result in protein reduction (Fig. S2). To test for specificity of the protein reduction, we also blotted for the IP_3_ receptor, which is highly abundant and can be detected on western blots using only a few oocytes or eggs. There was no reduction in IP_3_ receptor protein (Fig. 3A). SNAP23 depletion using Trim-away caused germinal vesicle breakdown (GVBD) in over 70% of follicle-enclosed oocytes after an ~18 hr culture period, whereas control oocytes that were either uninjected or that were injected with TRIM21/IgG did not undergo GVBD (Fig. 3B).

**Figure 3.**
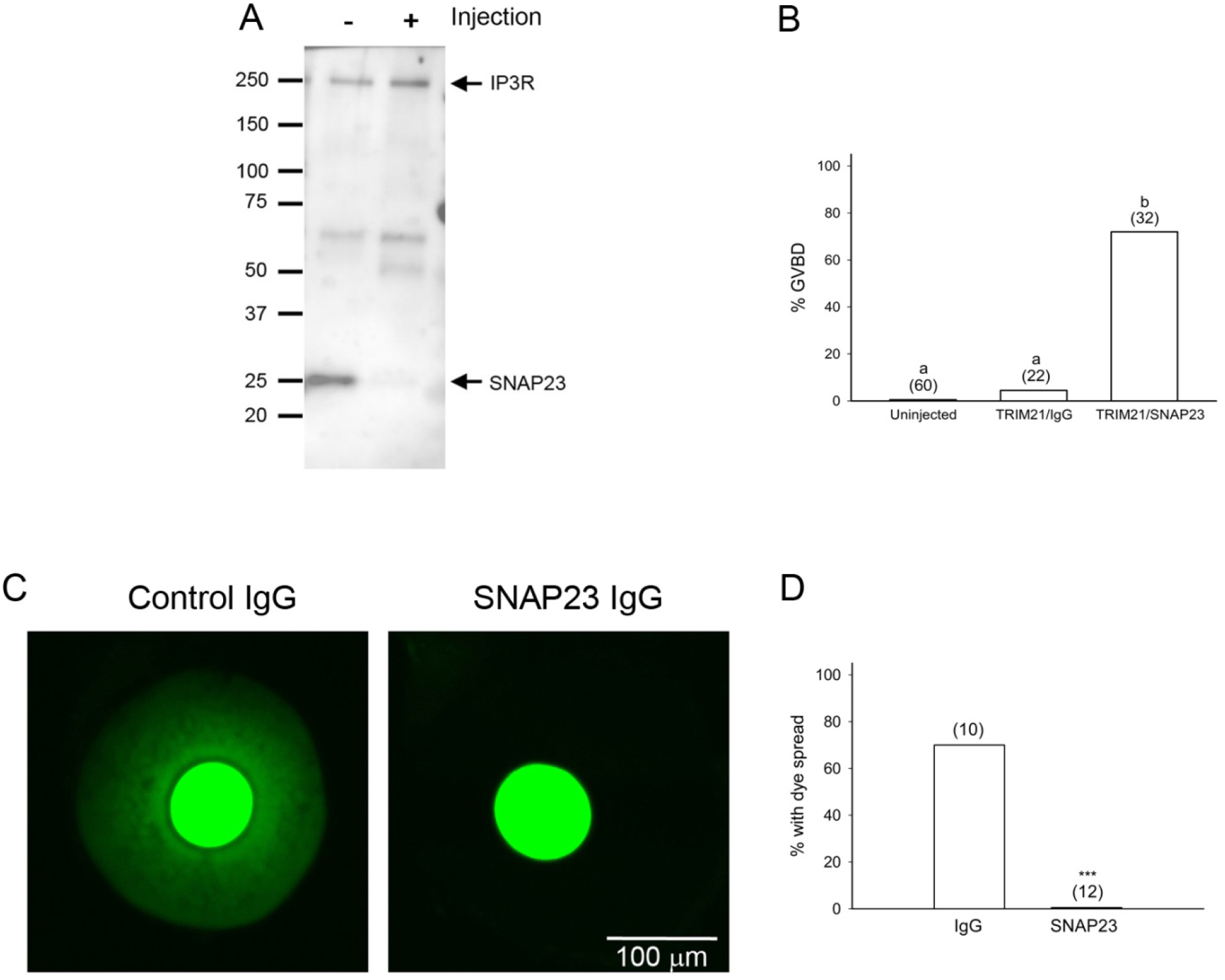
SNAP23 is required for maintenance of meiotic arrest and gap junction communication between the oocyte and cumulus cells. A) Western blot showing SNAP23 reduction using Trim-away, while levels of IP_3_ receptor were unaffected. B) SNAP23 depletion causes meiotic resumption in follicle-enclosed oocytes. The number of oocytes examined for each group is shown above each bar. Significance was determined using Fisher’s exact test and bars with different letters are significantly different. C) Alexa488 dye spreading from oocytes to follicle cells in control or SNAP23-depleted follicle-enclosed oocytes. D) Percentages of follicles that exhibited dye spreading. The number of follicles examined is stated above each bar. Statistical significance was determined using Student’s t-test. *** P<0.0001.

### SNAP23 depletion disrupts communication between the oocyte and somatic cells

Because SNAP23 depletion caused spontaneous maturation in follicle-enclosed oocytes, it is possible that inhibiting exocytosis disrupts communication between the oocyte and surrounding granulosa cells. To test this possibility, we first used Trim-away to deplete SNAP23 in follicle-enclosed oocytes, then injected oocytes with a low molecular weight fluorescent dye, Alexa488, that readily passes through gap junctions [40]. We used preantral follicles for these experiments due to the higher survival rate than we achieve using antral follicles. The Alexa488 dye spread throughout the granulosa cells in 70% (7/10) of control follicles (Fig. 3C,D). In contrast, the dye did not spread in any of 12 follicles in which SNAP23 was depleted (Fig. 3C,D).

The simplest explanation for the disconnection of gap junction communication between the oocyte and follicle cells when exocytosis is inhibited is that connexin 37 (Cx37), the protein that comprises gap junctions between the oocyte and cumulus cells [41], cannot be replaced in the plasma membrane following endocytosis. The half life of Cx37 is on the order of hours [42], so it needs to be continuously placed into the plasma membrane to form gap junctions. In the absence of gap junctions, the diffusion of cGMP from the cumulus cells to the oocyte would be stopped and this would activate PDE3A in the oocyte to lower cAMP levels to the point that the oocyte would no longer be able to maintain meiotic arrest [3, 43].

### Confirmation of the requirement of SNAP23 for maintenance of meiotic arrest, using a dominant negative approach

We used a dominant negative approach as a second way to inhibit SNAP23. We expressed a mutated form of SNAP25 (SNAP25Δ20), in which 20 amino acids in the C-terminus were deleted. This mutation disrupts the coiled coil domain in the C-terminus which is required for the formation of the SNARE complex [44], and inhibits exocytosis and stimulates meiotic resumption in *Xenopus* oocytes [44, 45]. Although SNAP25 was not identified in mouse oocytes, SNAP25Δ20 is likely to act as a dominant negative SNARE in oocytes because SNAP25 and SNAP23 can complex with the same t- and v-SNAREs [46, 47]. Both forms of SNAP25 were expressed in injected mouse oocytes (Fig. 4A). Expressing SNAP25Δ20 in follicle-enclosed oocytes robustly stimulated meiotic resumption (Fig. 4B), with nearly 100% of oocytes undergoing GVBD after an ~18 hr culture. Uninjected oocytes, or oocytes injected with SNAP25WT, did not resume meiosis during the culture period (Fig. 4B). These results using both Trim-away and a dominant negative SNARE support the hypothesis that SNAP23 is essential for constitutive exocytosis in mouse oocytes, and that constitutive exocytosis is needed to maintain meiotic arrest.

**Figure 4.**
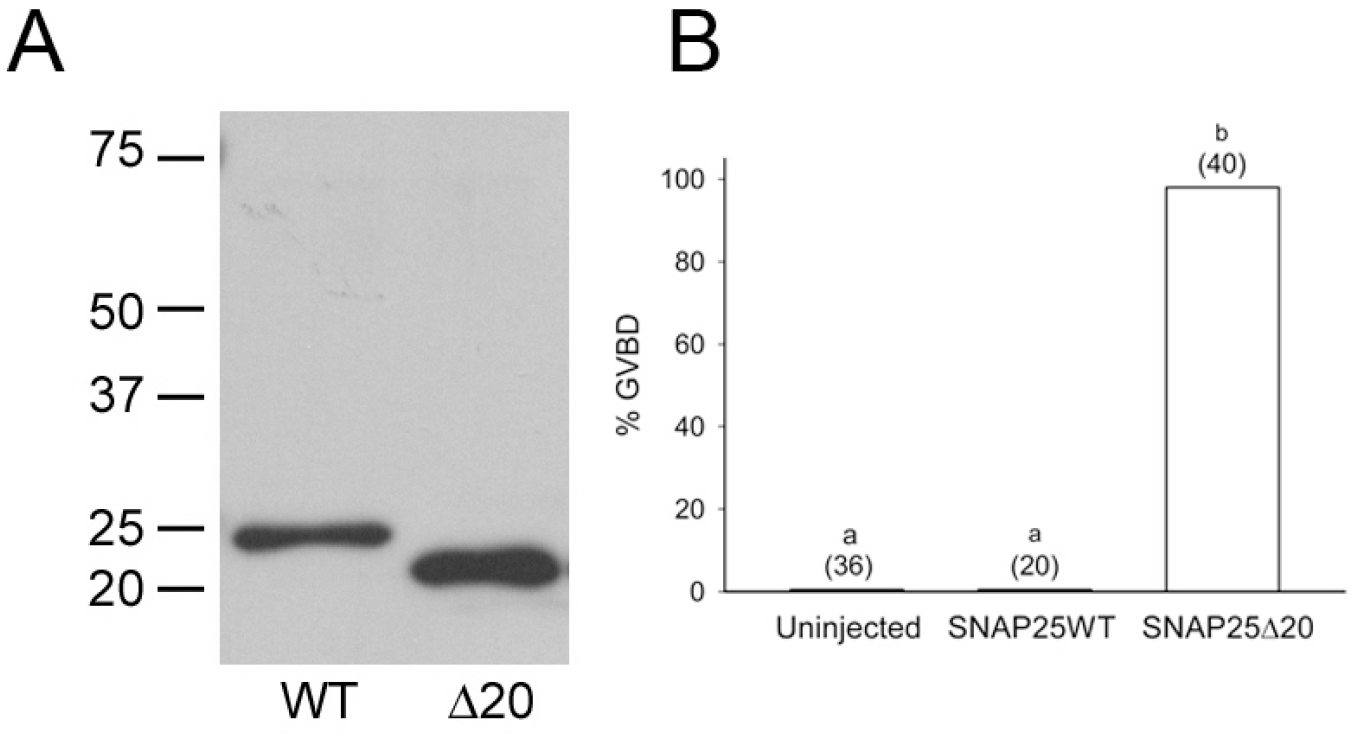
A dominant negative SNARE protein, SNAP25Δ20, stimulates meiotic resumption in follicle-enclosed oocytes. A) Both forms of SNAP25 were expressed in oocytes after microinjection. B) Percentage of follicle-enclosed oocytes that underwent GVBD following microinjection of SNAP25Δ20 or SNAP25WT. The number of oocytes examined is indicated above each bar. Significance was determined using Fisher’s exact test and bars with different letters are significantly different.

As discussed above, gap junctions are likely to be affected by depletion of SNAP23 and this probably inhibits the transfer of cGMP from the follicle cells to the oocyte, which causes meiotic resumption. Another possibility to account for meiotic resumption in the oocyte when exocytosis is blocked is that the oocyte cannot secrete factors such as GDF9 and BMP15 that act on the cumulus cells to maintain follicle integrity. This could include the continued synthesis of connexins on the follicle side. It is also possible that GPR3, the receptor needed for cAMP production in the oocyte [48–50], must be placed at the plasma membrane in order to signal to make cAMP, and in the absence of GPR3 at the surface the oocyte can no longer produce cAMP. Finally, it is possible that the oocyte secretes a ligand that stimulates GPR3 and in the absence of this ligand the oocyte can no longer produce cAMP. GPR3 is thought to be constitutively active and no ligand has yet been identified [51, 52], but there is still a possibility that the oocyte produces and secretes one.

### SNAP23 is required for regulated exocytosis

During maturation, oocytes gear up for the regulated exocytic event of cortical granule exocytosis. Because SNAP23 participates in constitutive and regulated exocytosis in other cell types [53, 54], we investigated if it is required for cortical granule exocytosis in mouse eggs using Trim-away. For these experiments, we co-injected immature oocytes with RNA encoding TRIM21 and an antibody against SNAP23 and matured the oocytes for ~18 hrs. Expression of TRIM21 and the SNAP23 antibody did not inhibit oocyte maturation and the mature eggs appeared healthy and formed first polar bodies. After maturation, we examined cortical granule exocytosis after providing a Ca^2+^ stimulus. We initiated Ca^2+^ release in eggs using the sulfhydryl reagent, thimerosal, which reliably produces a rapid series of long lasting, repetitive Ca^2+^ transients in mouse oocytes and eggs [35].

We detected cortical granule exocytosis following thimerosal treatment using two methods. In the first method, we removed the zonae pellucidae following injection of RNA and antibody, prior to overnight oocyte maturation. We then treated mature eggs with 100 μM thimerosal for 45 min and stained cortical granules with a fluorescent lectin [26]. Before thimerosal treatment, we detected cortical granules at the cortex opposite the meiotic spindle in untreated eggs in all 3 groups (Fig. 5A, left panels). The cortical granule-free domain was sometimes quite large, comprising around 50% of the oocyte cortex compared to the 20% cortical granule-free area observed in ovulated eggs [9]. This correlates with a recent study showing that cortical granule distribution and function in in vitro matured eggs is reduced compared to in vivo matured eggs [55]. Following thimerosal treatment, cortical granules were greatly reduced at the cortex and appeared to be more dispersed around the surface in small puncta of uninjected and control-injected eggs (Fig. 5A, right panels), demonstrating that under our conditions, cortical granule exocytosis occurs in in vitro matured eggs. In contrast, cortical granules were retained in the cortex of SNAP23 antibody-injected eggs, showing that exocytosis was blocked in these eggs (Fig. 5A).

**Figure 5.**
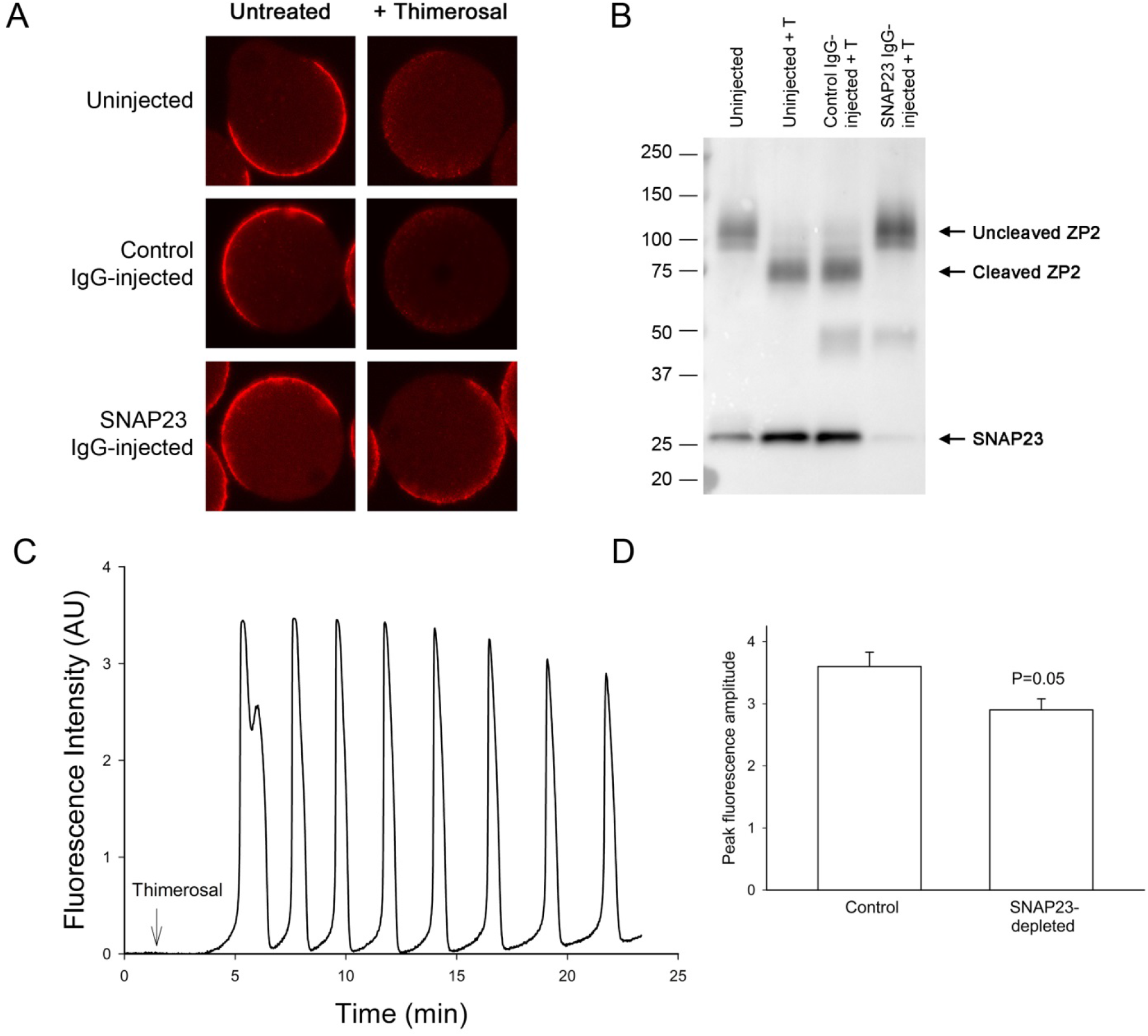
SNAP23 depletion inhibits cortical granule exocytosis. SNAP23 protein was degraded using Trim-away and cortical granules were labelled with TRITC-LCA after in vitro maturation and thimerosal treatment (A); or cortical granule exocytosis was assessed by ZP2 cleavage (B). The effectiveness of SNAP23 depletion is shown in (B). C) SNAP23 depletion does not inhibit Ca^2+^ release in response to thimerosal treatment. D) Peak amplitudes of Ca^2+^ release in SNAP23-depleted and control, uninjected eggs were not statistically significant (P=0.05).

In the second method, we left the zonae intact following microinjection and overnight culture. We treated eggs with thimerosal and examined ZP2 cleavage using western blot. As shown in Fig. 5B, uninjected eggs that were not treated with thimerosal contained the uncleaved form of ZP2. The band shifted following thimerosal treatment, demonstrating that ZP2 was cleaved. Likewise, ZP2 was cleaved in TRIM21-expressing, control IgG-injected eggs that were treated with thimerosal. ZP2 was not cleaved, however, in TRIM21-expressing, SNAP23 antibody-injected eggs treated with thimerosal. Probing the ZP2 blot with SNAP23 antibody showed that SNAP23 protein was barely detectable in antibody-injected eggs (Fig. 5B). SNAP23 depletion did not prevent Ca^2+^ release in response to thimerosal treatment (Fig. 5C), and the peak amplitudes were similar between SNAP23-depleted eggs and uninjected controls (Fig. 5D). This demonstrates that SNAP23 acts downstream of Ca^2+^ release. Taken together, these results show that SNAP23 is required for cortical granule exocytosis.

In conclusion, we show that SNAP23 is required for the maintenance of meiotic arrest and cortical granule exocytosis in mouse oocytes and eggs, respectively. We are currently working to identify other SNAREs that interact with SNAP23 to mediate constitutive and regulated exocytosis.

## ACKNOWLEDGMENTS

We thank Melina Schuh for providing helpful advice with Trim-away, Jurrien Dean for providing ZP2 antibody, Khaled Machaca for providing the SNAP25Δ20 and SNAP25WT constructs, Carmen Williams for providing the pIVT vector, Jim Watras for providing IP_3_ receptor antibody, and Laurinda Jaffe for helpful comments and suggestions.

## SUPPLEMENTAL FIGURES

**Figure S1.**
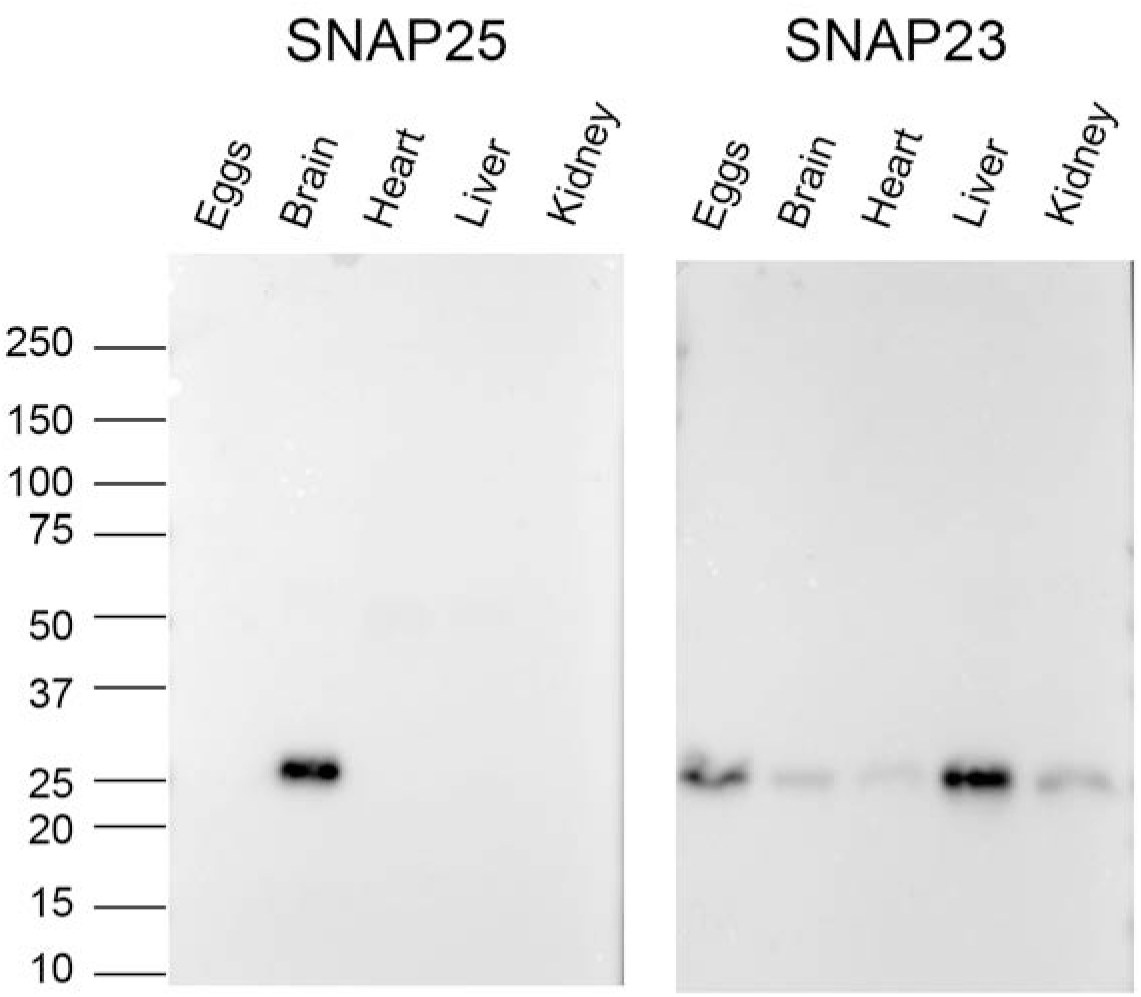
Expression of SNAP25 and SNAP23 in several tissues as well as in eggs. 1 μg of each indicated lysate was used per lane. SNAP25 was exclusively expressed in brain, whereas SNAP23 was expressed in each tissue that was examined.

**Figure S2.**
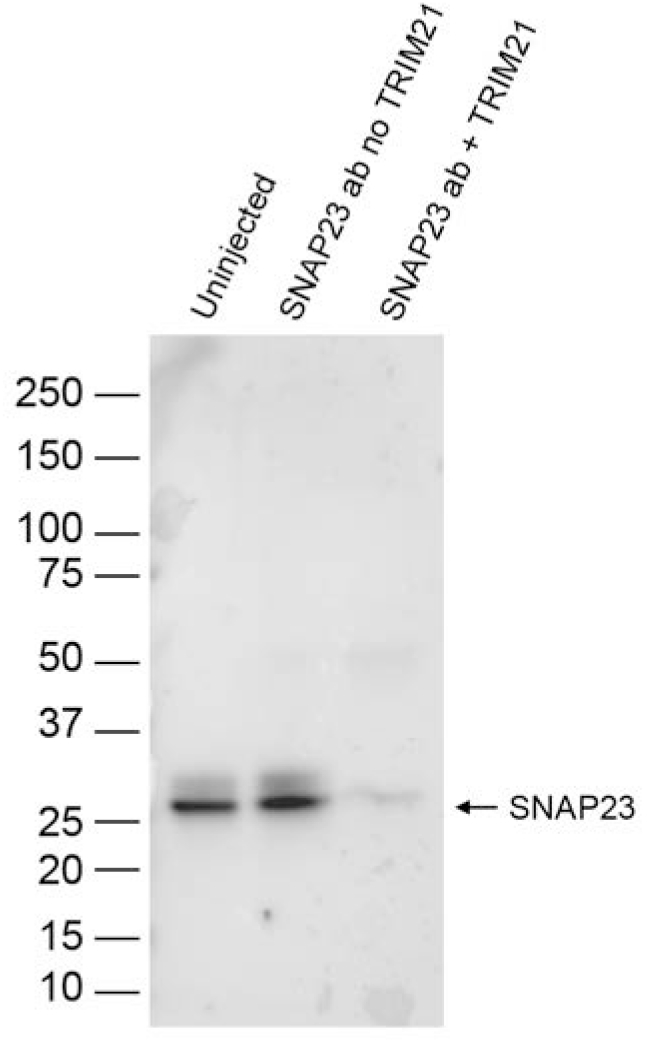
TRIM21 is required for SNAP23 depletion. Western blot showing expression of SNAP23 in oocytes that were injected with SNAP23 antibody with or without mRNA encoding TRIM21. Injected oocytes were matured overnight and lysates from 12 eggs were run in each lane.

